# Sensing their plasma membrane curvature allows migrating cells to circumvent obstacles

**DOI:** 10.1101/2021.03.26.437199

**Authors:** Ewa Sitarska, Silvia Dias Almeida, Marianne Sandvold Beckwith, Julian Stopp, Yannick Schwab, Michael Sixt, Anna Kreshuk, Anna Erzberger, Alba Diz-Muñoz

**Affiliations:** Cell Biology and Biophysics Unit, European Molecular Biology Laboratory, 69117 Heidelberg, Germany; Institute of Science and Technology Austria, 3400 Klosterneuburg, Austria

## Abstract

Cell migration is a hallmark out-of-equilibrium process in biology. In addition to persistent self-propelled motion, many cells display remarkable adaptive behaviors when they navigate complex environments within the body. Combining theory and experiments, we identify a curvature-sensing mechanism underlying obstacle avoidance in immune-like cells. The genetic perturbation of this machinery leads to a reduced capacity to evade obstructions combined with faster and more persistent cell migration in obstacle-free environments. We propose that the active polymerization of the actin cytoskeleton at the advancing edge of migrating cells is locally inhibited by the curvature-sensitive BAR protein Snx33 in regions with inward plasma membrane curvature. This coupling between actin and membrane dynamics leads to a mechanochemical instability that generates complex protrusive patterns at the cellular front. Adaptive motility thus arises from two simultaneous curvature-dependent effects, i) the specific reduction of propulsion in regions where external objects deform the plasma membrane and ii) the intrinsic patterning capacity due to the membrane-actin coupling that promotes spontaneous changes in the cell’s protrusions. Our results show how cells utilize actin- and plasma membrane biophysics to sense their environment, allowing them to adaptively decide if they should move ahead or turn away. On the basis of our findings, we propose that the natural diversity of BAR proteins may allow cells to tune their curvature sensing machinery to match the shape characteristics in their environment.

## Main text

The self-propelled motion of cells underlies many developmental, physiological, and pathological processes^1–3^. Former investigations revealed the rich physics of persistent cellular propulsion, clarifying for example how the non-equilibrium properties of a polar actin network can generate a steadily advancing leading edge^4–8^. In addition to persistent motion, migrating cells display remarkable adaptive behaviors. Cells within the body efficiently circumvent obstacles^9,10^ to navigate complex and dynamic environments^11^ but the physical basis of motility adaptation is not well understood. These adaptive behaviors are particularly important for cell types that encounter highly variable environments, such as immune cells and disseminating tumor cells. During migration, the outmost boundary of the cell – the plasma membrane – is deformed by the changing microenvironment. Thus, cells might decode their membrane curvature to decide if they should move ahead or turn away. Many migrating cells indeed express a variety of curvature-sensing proteins, which can directly interact with both the plasma membrane and the subjacent actin cytoskeleton^12^. Here we investigate the role of curvature sensing in adaptive motility.

The family of BAR domain-containing proteins could facilitate membrane curvature sensing. These proteins form crescent shaped membrane-binding dimers that can sense and generate curvature^12–14^, and are known to interact with regulators of the actin cytoskeleton through several auxiliary domains^12,15^. We identified candidates from this large protein family by measuring their expression profile in immune-like HL-60 cells before and after terminal differentiation, where they undergo substantial changes in gene expression and initiate rapid migration, resembling what occurs in the bone marrow *in vivo*^16^ (**Fig. 1a, b, Supplementary Fig. 1a**). The BAR domain protein Snx33 was 16-fold over-expressed in the curvature-rich state (**Supplementary Fig. 1b**). Based on its structure, Snx33 likely prefers comparatively small, positive (inward) curvatures (**Fig. 1c, Supplementary Fig. 2a-d**)^12^. We imaged its subcellular localization by confocal microscopy after fluorescent tagging and found that Snx33 is enriched in distinct membrane ruffles at the leading edge of these cells (**Fig. 1d, e**).

**Figure 1:**
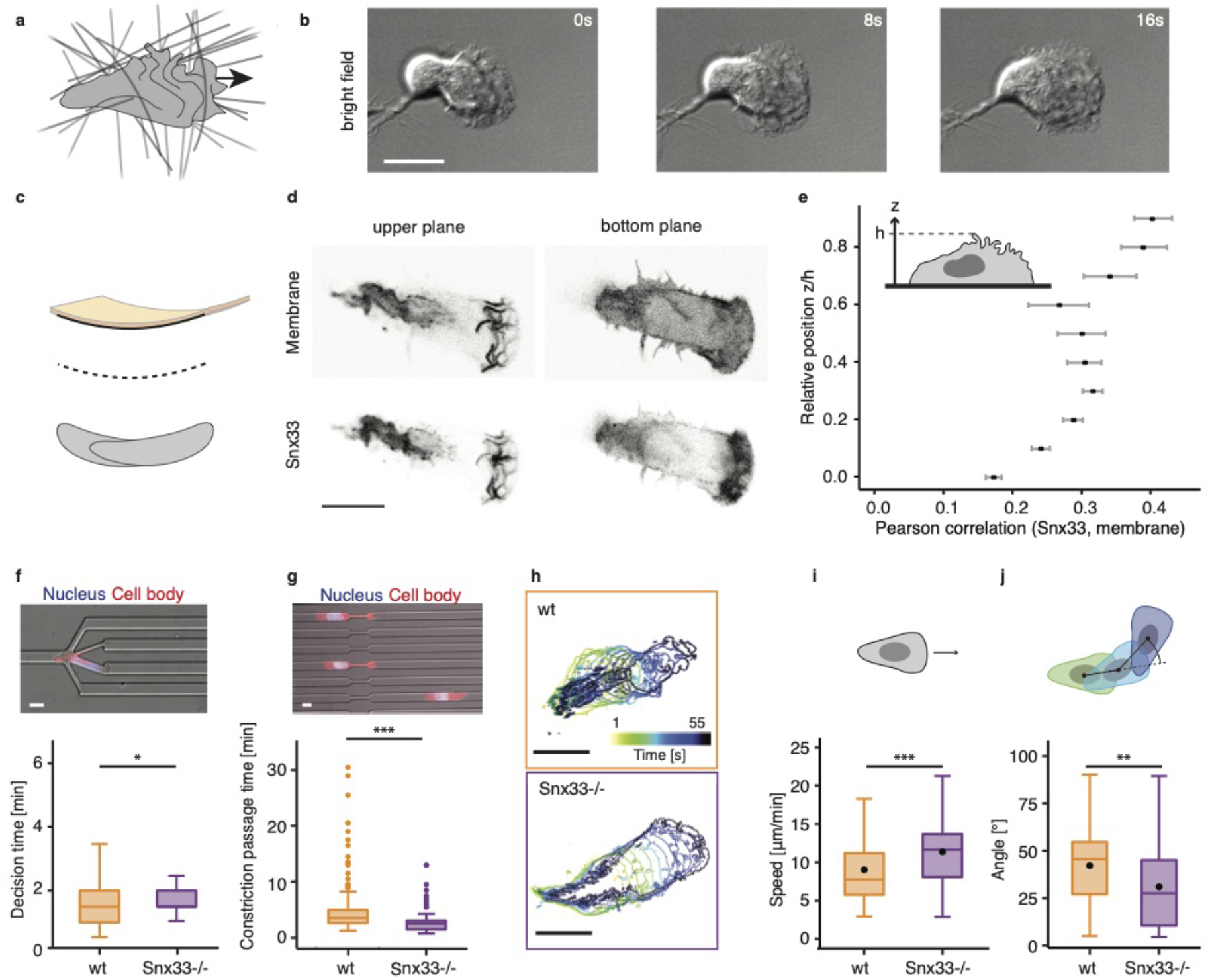
The curvature-sensing protein Snx33 is enriched in membrane waves and its loss impacts adaptive motility. **a)** The leading edge of migrating cells is characterized by intricate curvature patterns. Arrow indicates the direction of cell movement. **b)** Time-lapse bright-field imaging shows migrating immune-like differentiated HL-60 cells. **c)** The schematic illustrates the predicted binding preference for inward (positive) membrane curvature of Snx33, a BAR protein with one of the highest fold change during migration. **d)** Fluorescently-tagged Snx33 and membrane marker in upper and lower z-planes in a cell. **e)** Pearson correlation coefficient of fluorescently-tagged Snx33 and membrane marker at different heights acquired by confocal microscopy. n=10. Error bars denote standard error of the mean. **f)** Decision channel passage time (n=159 for wt, n=91 for Snx33-/-) in cells. **g)** Constriction passage time (n=234 for wt, n=158 for Snx33-/-) in cells. **h)** Cell body displacement over time in wt and Snx33-/- cells. **i)** Cell speed in wt and Snx33-/- cells and **j)** distribution of angles at which cells turn during migration. n=82 (wt), n=78 (Snx33-/-). Data from 3 independent biological replicates. Statistics: t-test or nonparametric Mann-Whitney-U-test. Scale bars = 10 μm. p<0.001 (***), p<0.01 (**), p<0.05 (*).

To address the role of curvature sensing in adaptive migration, we generated cells lacking Snx33 (Snx33-/-) using CRISPR/Cas9 genome editing (**Supplementary Fig. 3a, b**), and first assessed cell-scale migratory behavior. We positioned the cells in microfluidic devices^17,18^ where they migrated through channels with branching points with multiple differently sized pores. Snx33-/- cells required significantly more time than their wild-type counterparts to navigate these decision-points and find the path of least resistance (**Fig. 1f**, **Supplementary Fig. 4a, b**). However, in decision-free channels with or without a single constriction, Snx33-/- cells migrated faster than their wildtype counterparts (**Fig. 1g**, **Supplementary Fig. 4c**). Next, we quantified unconfined migration on planar substrates. We trained and used machine learning-based algorithms in ilastik^19^ to analyze movies of cells imaged by total internal reflection fluorescence microscopy (TIRFM) (**Fig. 1h**, see Methods for details). Similar as in decision-free channels, Snx33-/- cells migrated faster than wildtype cells (**Fig. 1i**). Moreover, while the motility of Snx33-/- cells was more persistent, wild-type cells were more prone to perform large spontaneous turns (**Fig. 1j**). Thus, the loss of Snx33 appears to render cells less effective in navigating a branching point, while more persistent in a decision-free environment, suggesting they have improved propulsion but lack the ability to adapt when faced with an inert obstacle.

Propulsion in immune-like cells depends on the active polymerization of actin at the leading edge of the cell^20,21^. Given that cell shape reflects changes in actin-rich protrusions^22^, we performed a quantitative and unbiased comparison of selected cell morphometric parameters, i.e., cell spreading, cell eccentricity and leading edge characteristics during decision-free migration on a planar substrate. As immune cells radically change their morphology in short periods of time, we averaged time-lapse movies to capture their dynamics. Snx33-/- cells spread to a larger extent, showed a more elongated morphology and bigger leading edge (**Fig. 2a-f, Supplementary Fig. 5a-e**); migration phenotypes that were independent of cell adhesion to the substrate (**Supplementary Fig. 5f-h**). Moreover, we measured the static tether pulling force of the plasma membrane with atomic force microscopy, and observed an increase from 61.58 pN for wildtype to 75.25 pN in Snx33-/- cells (which corresponds with an increase in membrane tension from 177.87 to 265.62 uN/m; see Methods for details; **Supplementary Fig. 6a, b**). Expressing fluorescently-tagged Snx33 reversed these changes in the morphology and mechanics of the leading edge thereby proving the specificity of the observed phenotypes (**Supplementary Fig. 5b, c and Supplementary Fig. 6b, c and d**). Notably, these phenotypes were not a consequence of defective differentiation, as the neutrophil differentiation marker CD11b was unperturbed in Snx33-/- cells (**Supplementary Fig. 6e**). Altogether, the observation that Snx33-/- cells have larger actin protrusions and higher membrane tension during decision-free migration indicates that deficient curvature sensing triggers an increase in actin polymerization at the leading edge even in the absence of obstacles.

**Figure 2:**
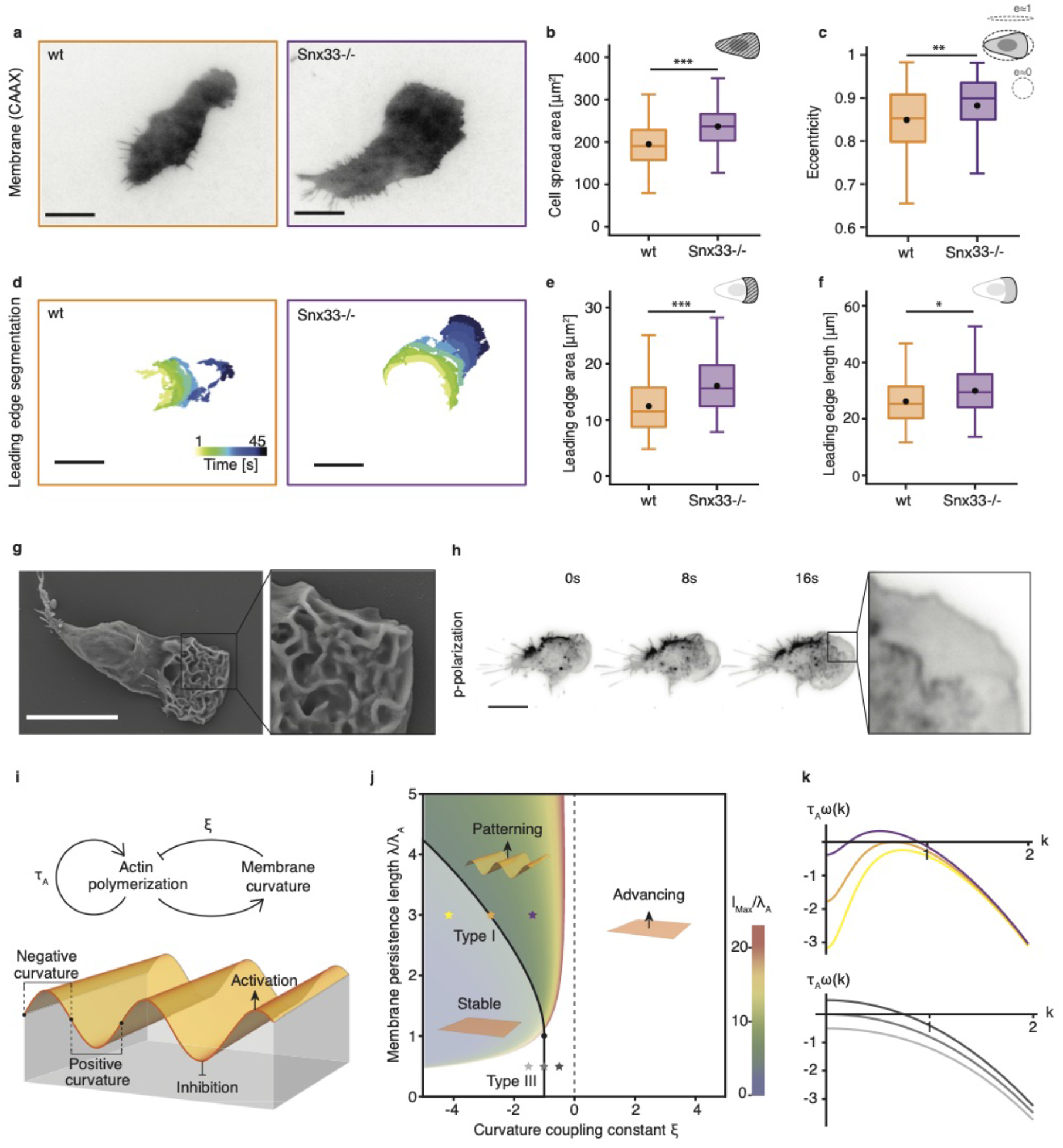
Genetic perturbation of Snx33-dependent curvature sensing alters cell and leading-edge morphology and may impact actin organization. **a)** TIRFM images of a wildtype and a Snx33-/- cell. **b)** Cell spread area and **c)** eccentricity differ between wildtype and Snx33-/- cells during movement. **d)** Leading-edge segmentation of a wildtype and a Snx33-/- cell. **e)** Leading-edge area and **f)** length differ between wildtype and Snx33-/- cells. n=82 (wt), n=78 (Snx33-/-). **g)** Scanning electron microscopy (SEM) image of a wildtype cell with zoom-in at the leading-edge. **h)** Time-lapse p-polarization of p-TIRFM imaging reveals dynamic membrane waves at the leading-edge. **i,j)** Negative coupling between membrane curvature and actin activity leads to spontaneous patterning. The transition from decaying to growing patterns corresponds to a type-I instability at the critical curvature coupling constant 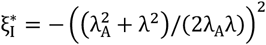 for membrane persistence lengths λ > λ_A_. Color scale denotes the wavelength with the highest growth rate l_Max_. **k**) Dispersion relations near the critical coupling constants for λ > λ_A_ (type-I) and λ < λ_A_ (type-III) at the positions marked in **j**). Statistics: t-test or non-parametric Mann-Whitney-U-test. Scale bars = 10 μm. p<0.001 (***), p<0.01 (**), p<0.05 (*).

These findings suggest that Snx33 mediates an intrinsic coupling between actin organization and membrane curvature. Such feedback effects have been proposed to trigger mechanochemical instabilities^23,24^ indicative of an intrinsic patterning capacity. Motile cells often feature wave-like patterns of actin activity and plasma membrane ruffles at their leading edge^20,21,25^ as observed by scanning electron microscopy (SEM) and polarized total internal reflection fluorescence microscopy (p-TIRFM), (**Fig. 2g, h and Supplementary Fig. 7a, b**). To link the dynamic membrane-actin patterns to the putative curvature-sensing function of Snx33, we analyzed the mechanochemical instability arising from a corresponding coupling between actin-dependent force generation and membrane curvature with a minimal set of equations and parameters (**Fig. 2i-k and Supplementary Note**). We considered a Helfrich membrane with a bending rigidity κ and surface tension γ ^26,27^ subject to an active pressure field due to actin polymerization. For the simplified case of a 1D pattern on a periodic domain x, we obtain the following equation for the membrane curvature C

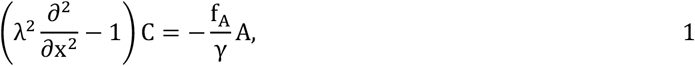

in which 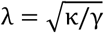 is the membrane persistence length, A is the concentration field of a generic activator of actin polymerization, and f_A_ is the force per activator molecule. The activator concentration is taken to evolve according to a reaction-diffusion equation, in which we consider self-activation of A ^20^ on a timescale τ_A_ and a curvature-dependent regulatory term r_C_(C)

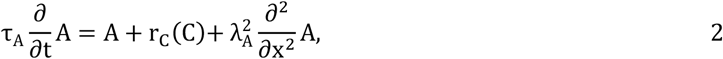

in which 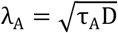 is a reaction-diffusion length scale with diffusion coefficient D. For regulation in the regime of small curvatures, the second reaction term can be truncated beyond linear order and written as r_C_(C) = ξγ/f_A_ C with a dimensionless coupling constant ξ. Linear stability analysis of Eqs. 1–2 then yields the dispersion relation for the growth rate ω(k) of the perturbation mode with wavenumber k

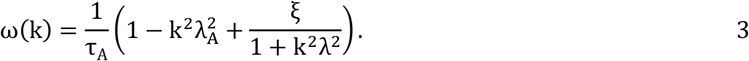

The parameter region in which spontaneous patterning occurs (where ω_Max_ > 0 with k_Max_ > 0) is shown in **Fig. 2j.** Spontaneous patterns emerge in this system only when there is a *negative* coupling between curvature and actin activity, such that actin polymerization decreases in regions of inward (positive) plasma membrane curvature (**Fig. 2g, h and i**), distinct from previously considered scenarios with curved activators^28–30^. Thus, we find that the local down-regulation of actin polymerization in regions with positive membrane curvature can generate mechanochemical patterns at the leading edge of migrating cells (**Fig. 2g, h and Supplementary Fig. 7a, b**).

This theoretical result agrees with the likely binding preference of Snx33 to membrane regions with small positive curvatures (**Supplementary Fig. 2a-d**) and is compatible with the observation that actin polymerization appears increased in Snx33-/- cells **(Fig. 2d, e, Supplementary Fig. 5c, Supplementary Fig. 6b).** But how does Snx33 inhibit actin polymerization? Several BAR domain proteins, including Snx33, directly bind and regulate the activity of actin nucleator promoting factors from the WASP family^12,31^. WAVE2 is the main member of this family in neutrophils and earned its name because it localizes in a wave pattern during cell migration^20^. What regulates WAVE2 binding to the membrane, and whether and how that determines ruffle topography remains poorly understood^32^. Our analysis of the mechanochemical feedback between membrane curvature and an actin activator such as WAVE2 (Eqs. 1–2) suggests that the dominant wavelength of the pattern should increase in response to a decrease in the magnitude of the curvature coupling coefficient ξ (**Fig. 3a**). To test this prediction, we imaged WAVE2 patterns in Snx33-/- and wild-type cells during migration by TIRFM (**Fig. 3b-d**). Snx33-/- cells showed an increase in both the length and the width of WAVE2 patches (**Fig. 3e, Supplementary Fig. 7c, d**). A mechanochemical curvature coupling would furthermore imply that the change in WAVE2 patch morphology should accompany a longer wavelength of the curved ruffles on the plasma membrane (**Fig. 3a**). To test this, we quantified the effective ruffle wavelength from SEM images and indeed observed a significantly less tight arrangement of ruffles for Snx33-/- cells when compared with their wild-type counterparts (**Fig. 3f, g, Supplementary Fig. 7e, f**). Notably, to date, only very drastic WAVE2 patch phenotypes have been reported (complete abrogation or increase in number), which lead to a severely disrupted morphology and capacity to migrate^33,34^. The milder perturbation introduced by the genetic disruption of curvature sensing by Snx33 allowed the identification of the key molecular circuit that couples the active mechanics of the actin network with the biophysics of the membrane to generate complex patterns at the leading edge of migrating cells.

**Figure 3:**
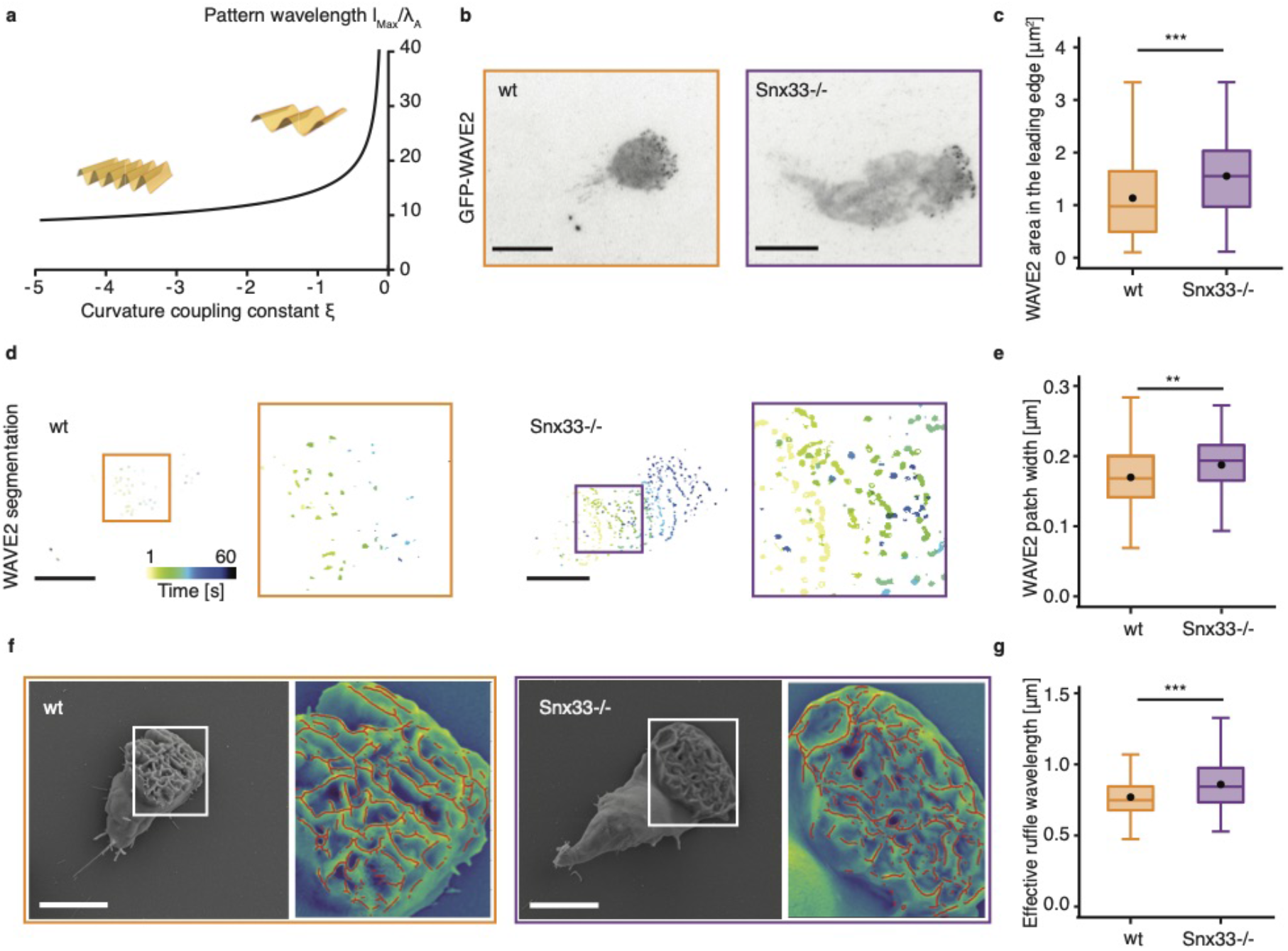
Disruption of curvature sensing alters actin activator patterns and membrane ruffles. **a)** Our theoretical calculation predicts that the dominant wavelength l_Max_ of the mechanochemical pattern should increase as the magnitude of the curvature coupling ξ decreases (here shown for λ/λ_A_ = 4). **b)** TIRFM images of a wildtype and Snx33-/- cell show the distribution of the actin activator WAVE2. **c)** The total area occupied by WAVE2 patches in the leading edge increases upon perturbation of Snx33. **d)** Segmentation of dynamic WAVE2 patches in a wildtype and Snx33-/- cell. **e)** The width of WAVE2 patches increases upon perturbation of Snx33. n=82 (wt), n=78 (Snx33-/-). **f)** SEM images with ruffle segmentations (red) show the leading edges of a wildtype and Snx33-/- cells from the 50^th^ percentile of the distribution shown in panel g). **g)** The effective ruffle wavelength increases upon perturbation of Snx33. n=175 (wt), n=170 (Snx33-/-). Statistics: t-test or nonparametric Mann-Whitney-U-test. Scale bars = 10 μm. p<0.001 (***), p<0.01 (**), p<0.05 (*).

This intrinsic patterning capacity can facilitate adaptive motility by reducing persistence and promoting the random evasive maneuvers we observed (**Fig 1j**). But the particular form of Snx33-mediated membrane-actin coupling suggests an additional more direct effect, in which propulsion is reduced specifically where the presence of an obstacle in the path of the cell is likely, i.e. where external indentations generate an increased fraction of regions with positive curvature. To gain further insights into the object-avoidance response of Snx33-deficient cells, we devised a reductionistic assay to investigate how cells steer away from a cellular obstacle, a common phenomenon to many cell types also known as contact inhibition of locomotion (CIL)^20,35^. We seeded a higher density of cells and imaged their 2D migration in the presence of cell-cell interactions by TIRFM. Snx33-/- cells collided more often and formed larger cell-cell contacts compared to wild-type cells (**Fig. 4a, b)**. To gain molecular insight into the role of Snx33 in CIL, we simultaneously live-imaged the WAVE2 complex and Snx33 at the leading edge of freely moving cells. While both proteins largely colocalized in most of the cell, they were anticorrelated in the highly negatively (outward) curved leading edge (**Supplementary Fig. 8a, b**). Here, Snx33 levels decreased while WAVE2 accumulated (**Supplementary Fig. 8c, d**), consistent with Snx33 binding positive curvature and restricting WAVE2 to negatively curved areas. Moreover, when cells collided, Snx33 localized to the cell contact area that is largely devoid of outward curvature, where subsequently WAVE2 disappeared and relocated to contact free zones of the plasma membrane. This resulted in cell repolarization by formation of a new leading edge (**Fig. 4c, d**). To assess whether the dysfunctional CIL response in Snx33-/- cells is indeed due to impaired WAVE2 inhibition at the contact area, we followed WAVE2 localization before and after collision in Snx33-/- cells. In contrast to wildtype cells, Snx33-/- cells indeed failed to remove WAVE2 from the contact site (**Fig. 4e, Supplementary Fig. 8e**). Altogether, we show a key role for membrane topography and the curvature sensing protein Snx33 in regulating actin polymerization during CIL. Specifically, Snx33 restricts WAVE to negatively curved areas and thus leads to cell repolarization towards a contact free zone upon collision with an obstacle.

**Figure 4:**
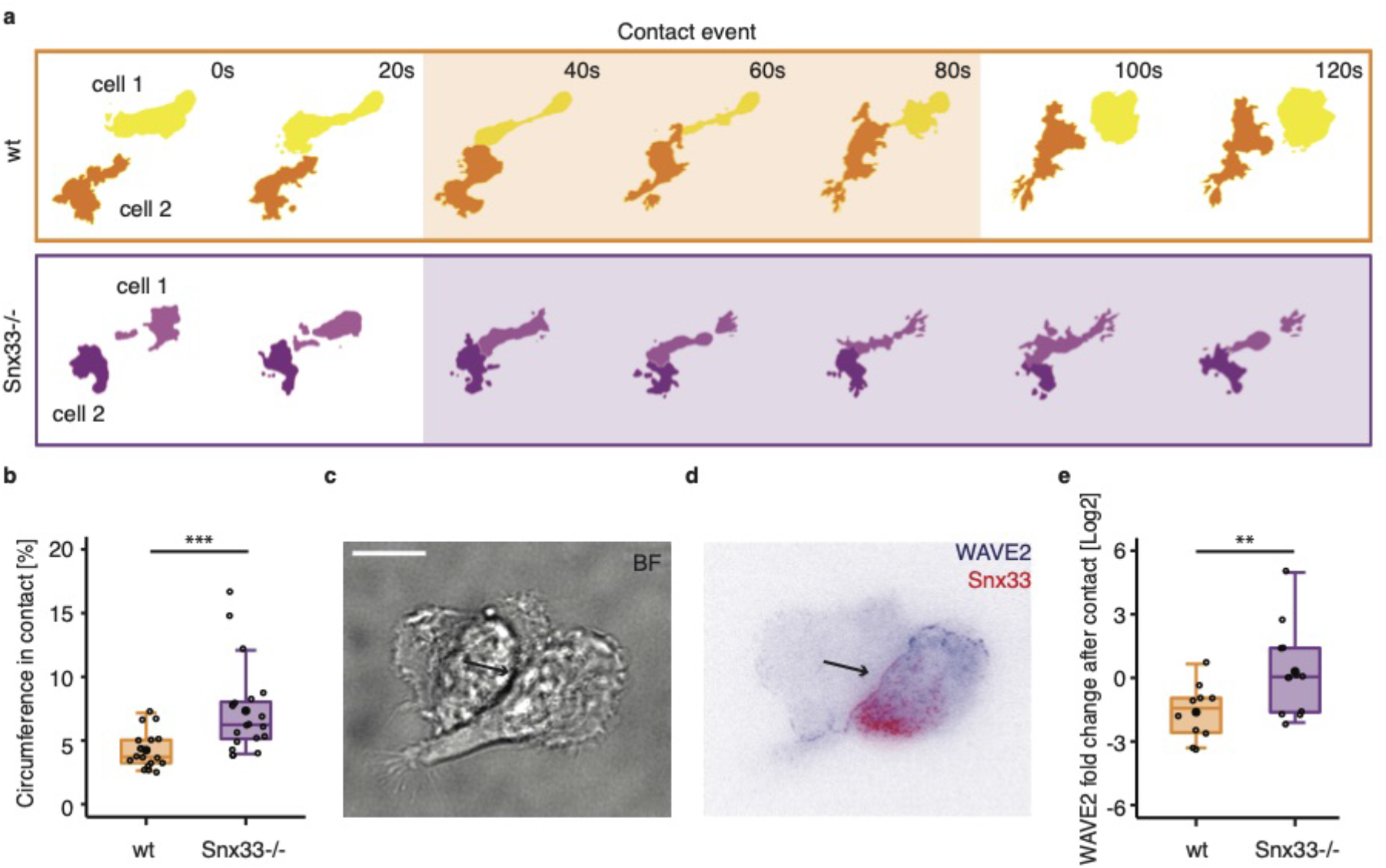
Snx33 regulates contact avoidance by locally inhibiting the WAVE2 complex. **a)** Segmentation of a contact event between two wildtype (upper panel) and two Snx33-/- (lower panel) cells. Shaded area highlights the duration of contact. **b)** Percentage of cell circumference in contact with another cell in wildtype (n=18) and Snx33-/- (n=20) cells. **c)** Bright-field and **d)** TIRFM imaging of a cell-cell contact in wildtype cells with fluorescently-tagged Snx33 and a component of WAVE2. Arrow is pointing towards cell-cell contact. **e)** WAVE2 fold change after cellcell contact in wildtype (n=9) and Snx33-/- (n=10) cells. Statistics: t-test or non-parametric Mann-Whitney-U-test. Scale bars = 10 μm. p<0.001 (***), p<0.01 (**), p<0.05 (*).

Object avoidance is key not only for the migration of immune and cancer cells in complex tissue environments, but also fundamental during embryogenesis and collective migration *in vivo*^36^. Combining theory, genetic perturbations, microscopy and microfluidics we identified a local mechanochemical circuit underlying adaptive motility in immune-like cells. We show that by coupling plasma membrane curvature with WAVE2-driven actin polymerization, the curvature sensing protein Snx33 facilitates adaptive motility in two distinct and complementary ways. On the one hand, the mechanochemical feedback between actin and curvature results in an intrinsic patterning capacity which destabilizes the leading edge and reduces migration persistence. The ability to spontaneously reorient and explore the microenvironment with dynamic protrusions in and of itself contributes to more efficient navigation in crowded contexts^17,33,37–40^. On the other hand, while a diverse range of feedback mechanisms can conceivably lead to shape instabilities^41–44^, Snx33 implements the coupling between membrane curvature and actin activity in a way that is uniquely suited to aid obstacle avoidance. Locally reducing propulsion in regions of inward curvature specifically steers the cell away from the direction in which the presence of an obstacle on the outside is most likely. Thus, Snx33 and WAVE2 form a non-genetic adaptive circuit that decodes the cell’s local membrane topography and induces a response to decide if it should move ahead or turn away.

The machinery we identified morphologically solves the object avoidance problem by exploiting the physics of the materials it is made of, specifically the biophysics of the plasma membrane and the active properties of the subjacent actin network. Given these findings and the diversity of BAR domain proteins present in cells, it is conceivable that on long timescales, cells tune their expression levels to match the curvature statistics of their microenvironment, implying that a cell’s curvature-sensing machinery may itself constitute a structural prediction of its surroundings.

---

**Supplementary Figure 1:**
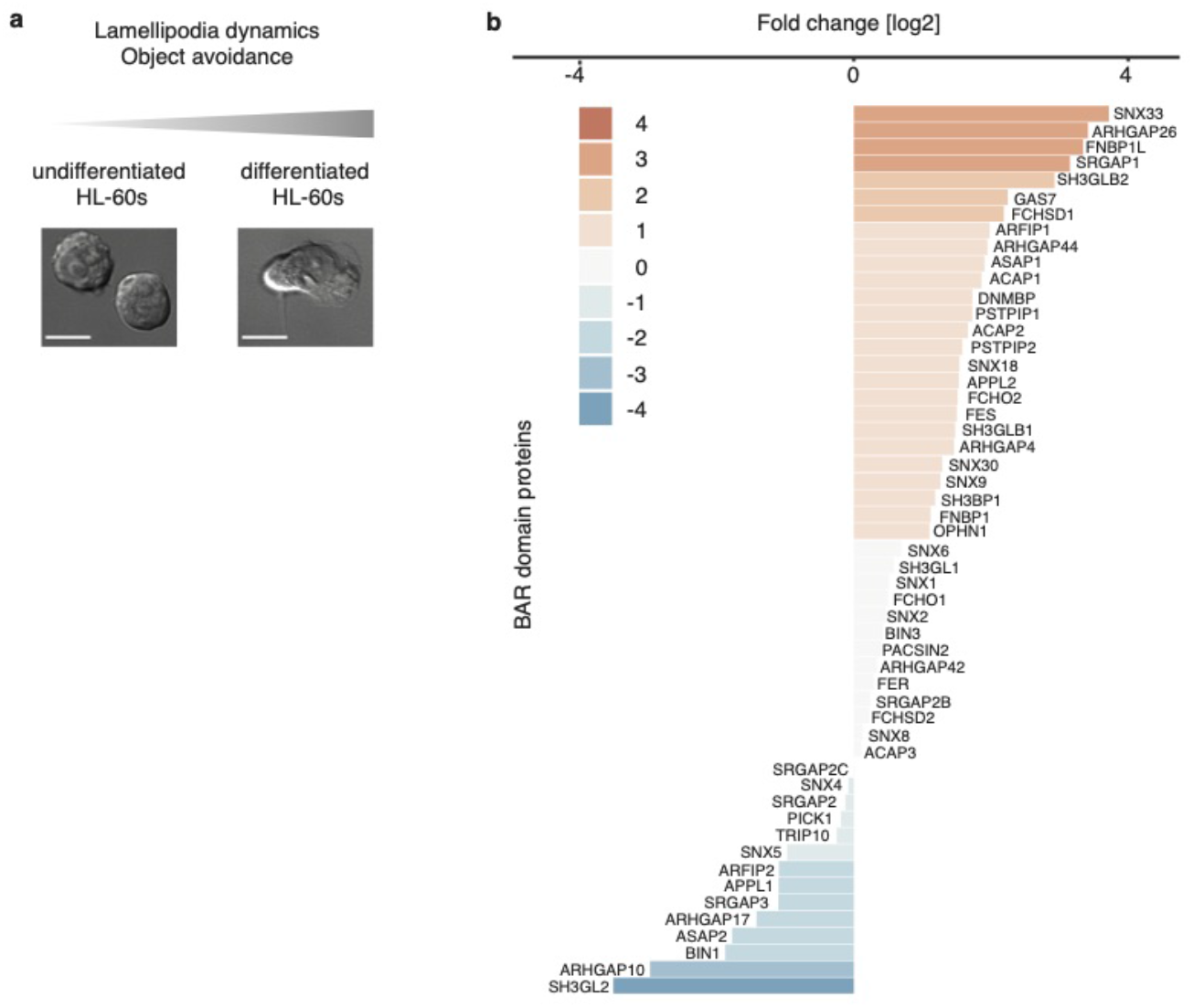
HL60 differentiation in vitro and fold change in expression of BAR domain proteins. **a)** Schematic illustrating changes in HL-60 cells during its differentiation. **b)** Over-expressed (orange) and under-expressed (blue) BAR domain genes between undifferentiated (non-migratory) and differentiated (migratory) HL-60 cells. Scale bars = 10 μm.

**Supplementary Figure 2:**
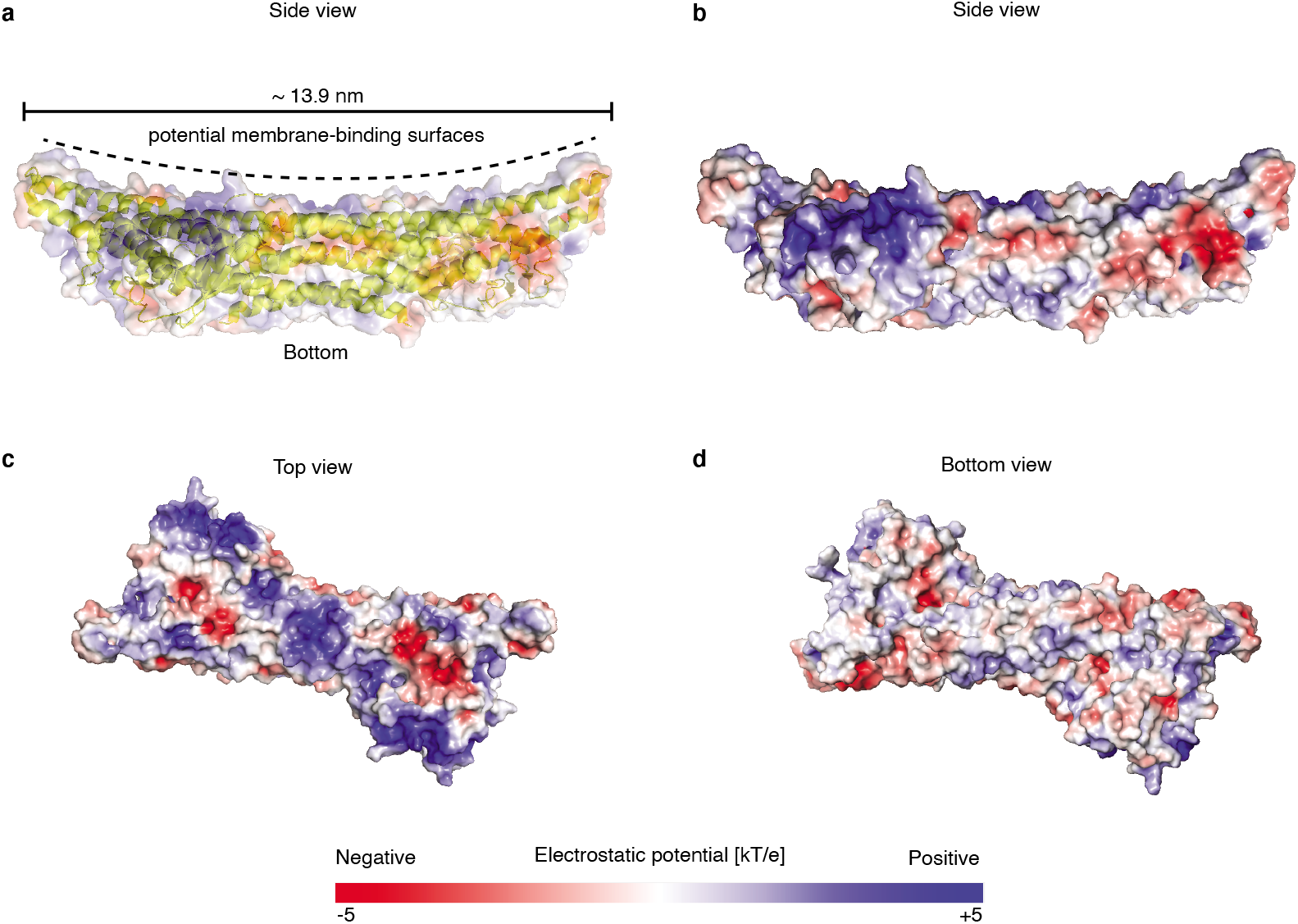
Structure of Snx33 membrane binding unit (PX-BAR) strongly suggests binding to inward (positive) membrane curvature. **a)** Structure of PX-BAR domains of Snx33 (4AKV) together with electrostatic surface representation (side view). Dotted line indicates the potential membrane-binding surfaces, while solid line shows overall dimensions of the dimer. **b-d)** Electrostatic surface representation of 4AKV side view **(b)**, top view **(c)** and bottom view **(d)**. Visualizations were created in PyMOL (The PyMOL Molecular Graphics System, Version 2.4.2 Schrödinger, LLC).

**Supplementary Figure 3:**
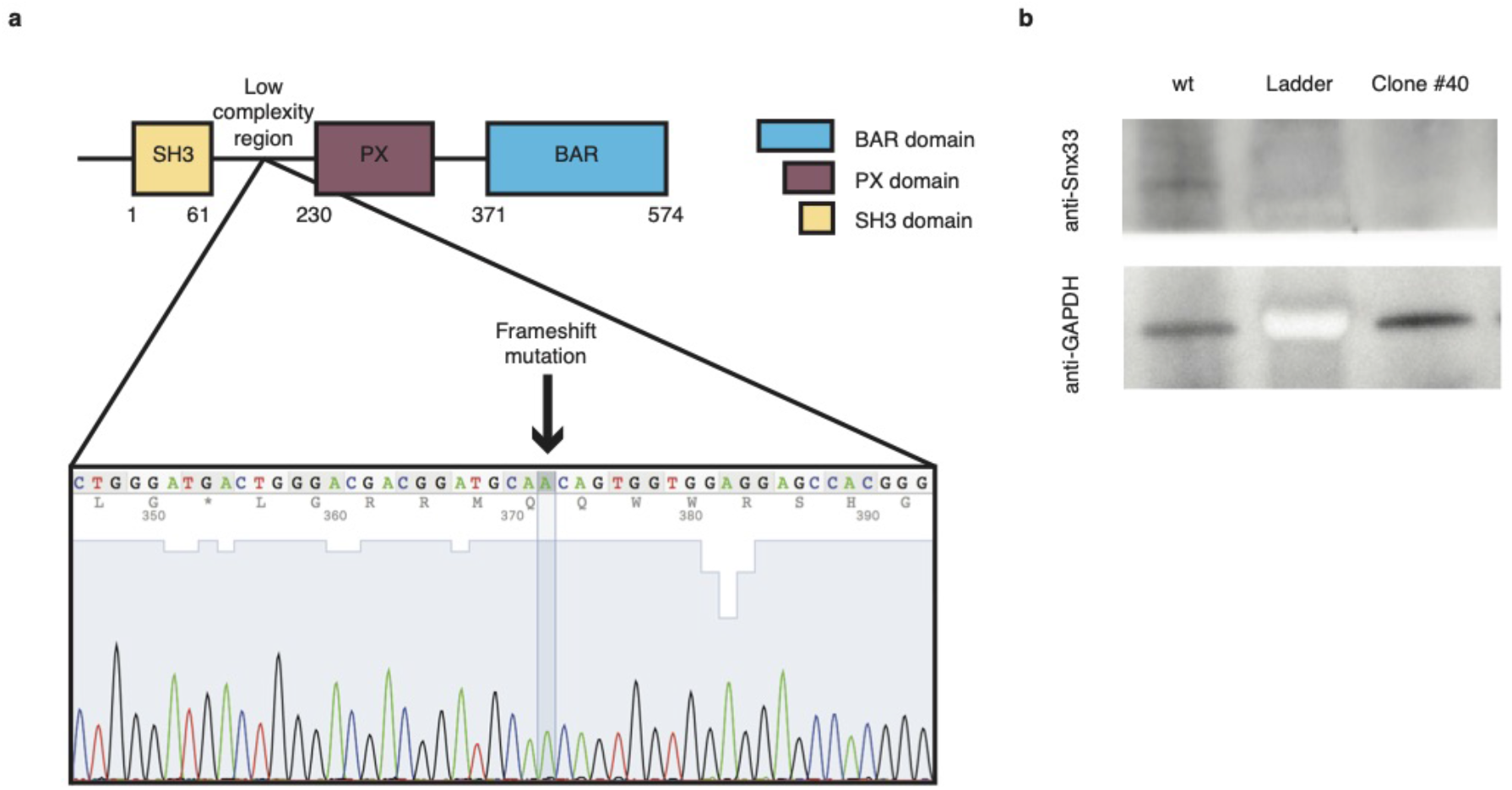
Snx33 knockout cell line validation. **a)** Snx33 clone sequencing confirming a frameshift mutation. **b)** Snx33 and GAPDH Western blots of wt cells and Snx33-/- clone.

**Supplementary Figure 4:**
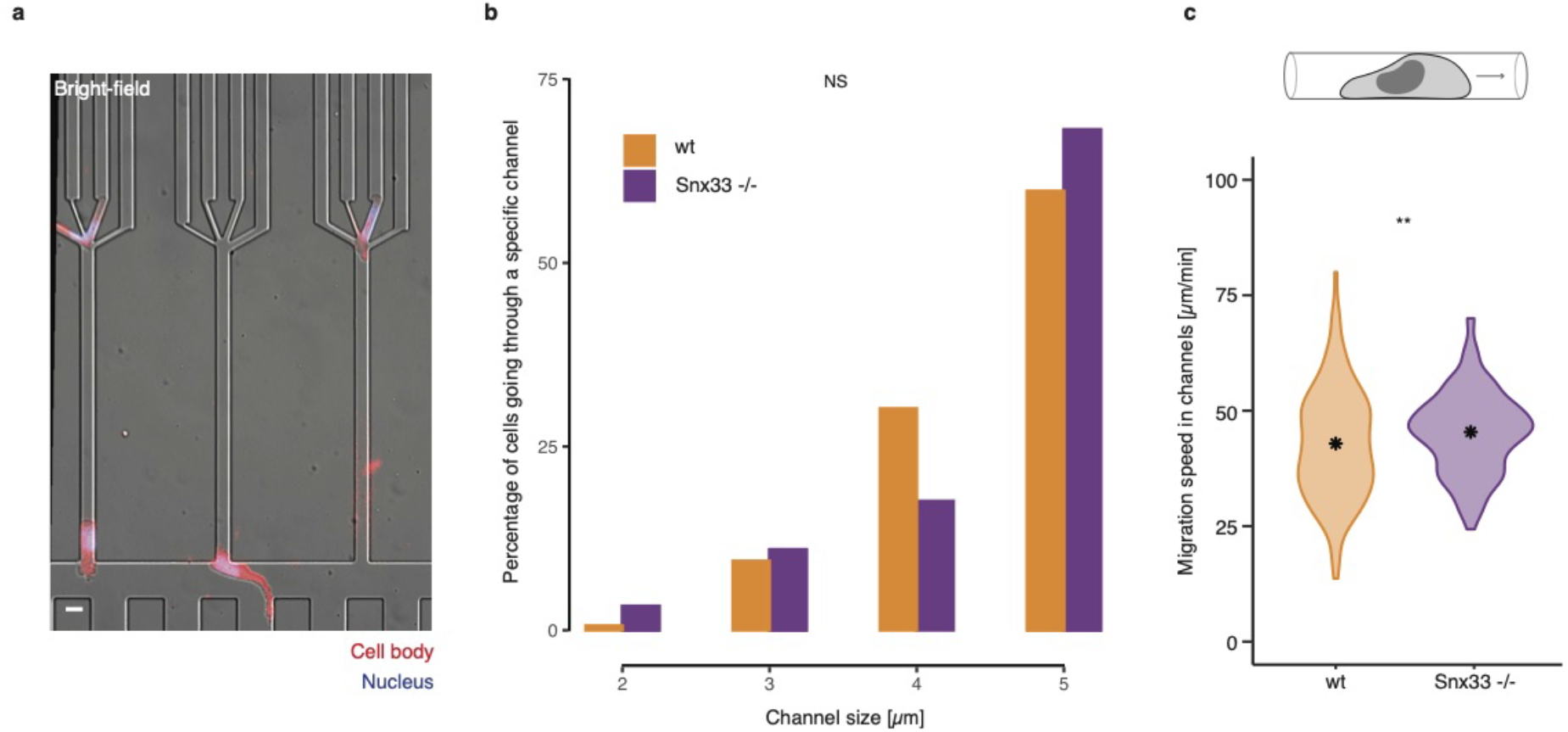
Snx33 knockout cells are faster and choose the path of the least resistance. **a)** Overlay of bright-field, nuclei and cell body images of cells migrating in PDMS-based devices with a decision point. **b)** Frequency of wt and Snx33-/- cells choosing a channel of a certain size (n = 159 for wt, n = 91 for Snx33-/-). **c)** Migration speed in straight channels (n=235 for wt, n=169 for Snx33-/-). Data from 3 independent biological replicates. Scale bar = 10 *μ*m. p<0.001 (***), p<0.01 (**), p<0.05 (*).

**Supplementary Figure 5:**
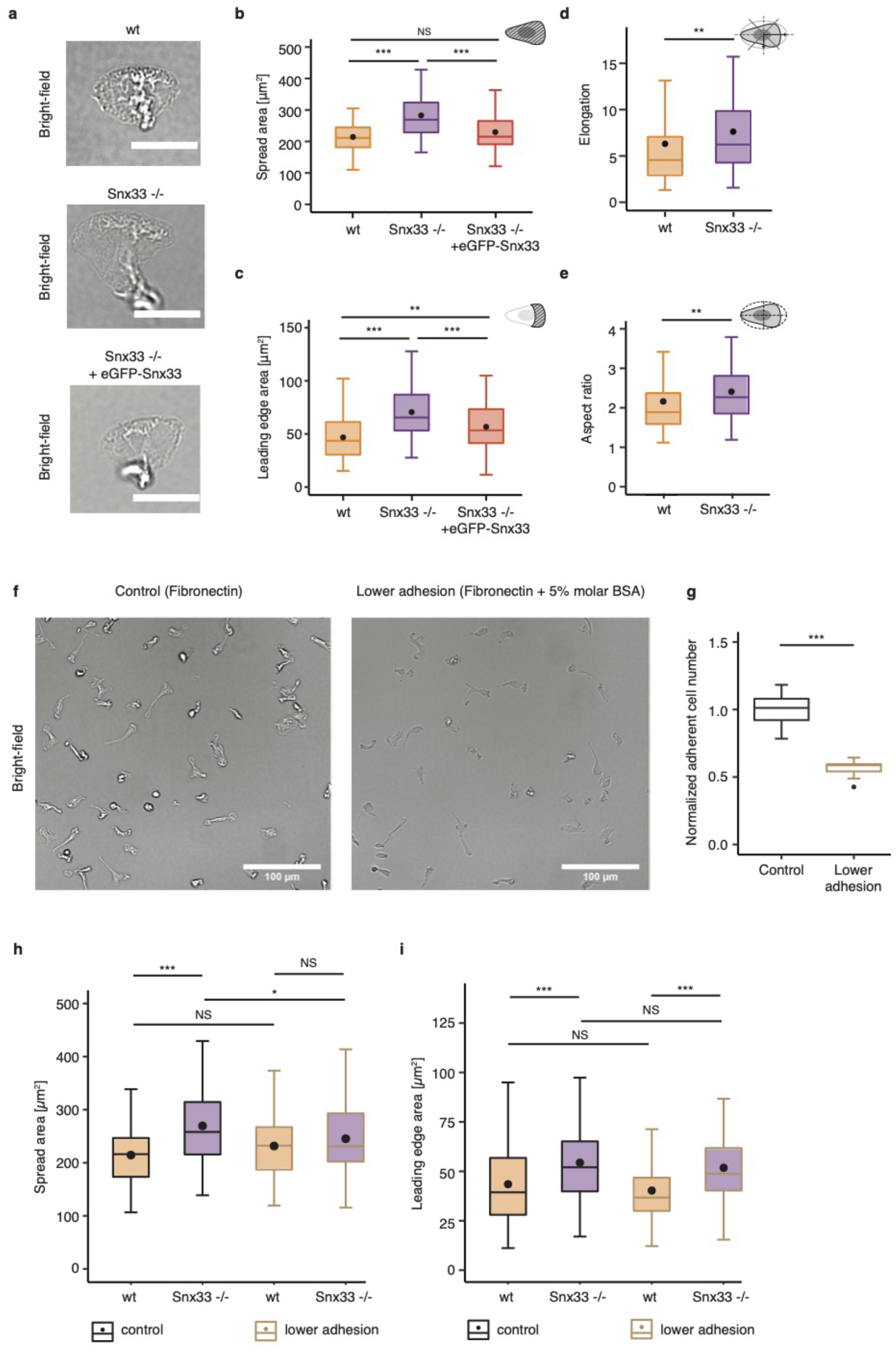
Cell and leading-edge morphology in Snx33 knockout and wt cells. **a)** Example images of bright-field of wild type, Snx33-/- and Snx33-/- with eGFP-tagged Snx33 cells. Quantification of **b)** cell spread and **c)** leading edge area. Data from 3 independent experiments. n= 67 (wt), n=73 (Snx33-/-), n=67 (Snx33-/- +GFP-Snx33) **d)** Quantification of cell elongation and **e)** aspect ratio from TIRFM images. n=82 (wt), n=78 (Snx33-/-). **f)** Example bright-field images of wt cells on control (fibronectin only) and lower adhesion substrate (fibronectin with 5% molar BSA). **g)** Adhesion-dependent cell number quantification. Data from two independent experiments. Quantification of **h)** cell and **i)** leading edge area. n = 82 (wt), n = 102 (Snx33-/-); Lower adhesion: n= 53 (wt), n = 72 (Snx33-/-). Statistics: t-test and Mann-Whitney-U-Test. Scale bars = 10 μm. p<0.001 (***), p<0.01 (**), p<0.05 (*).

**Supplementary Figure 6:**
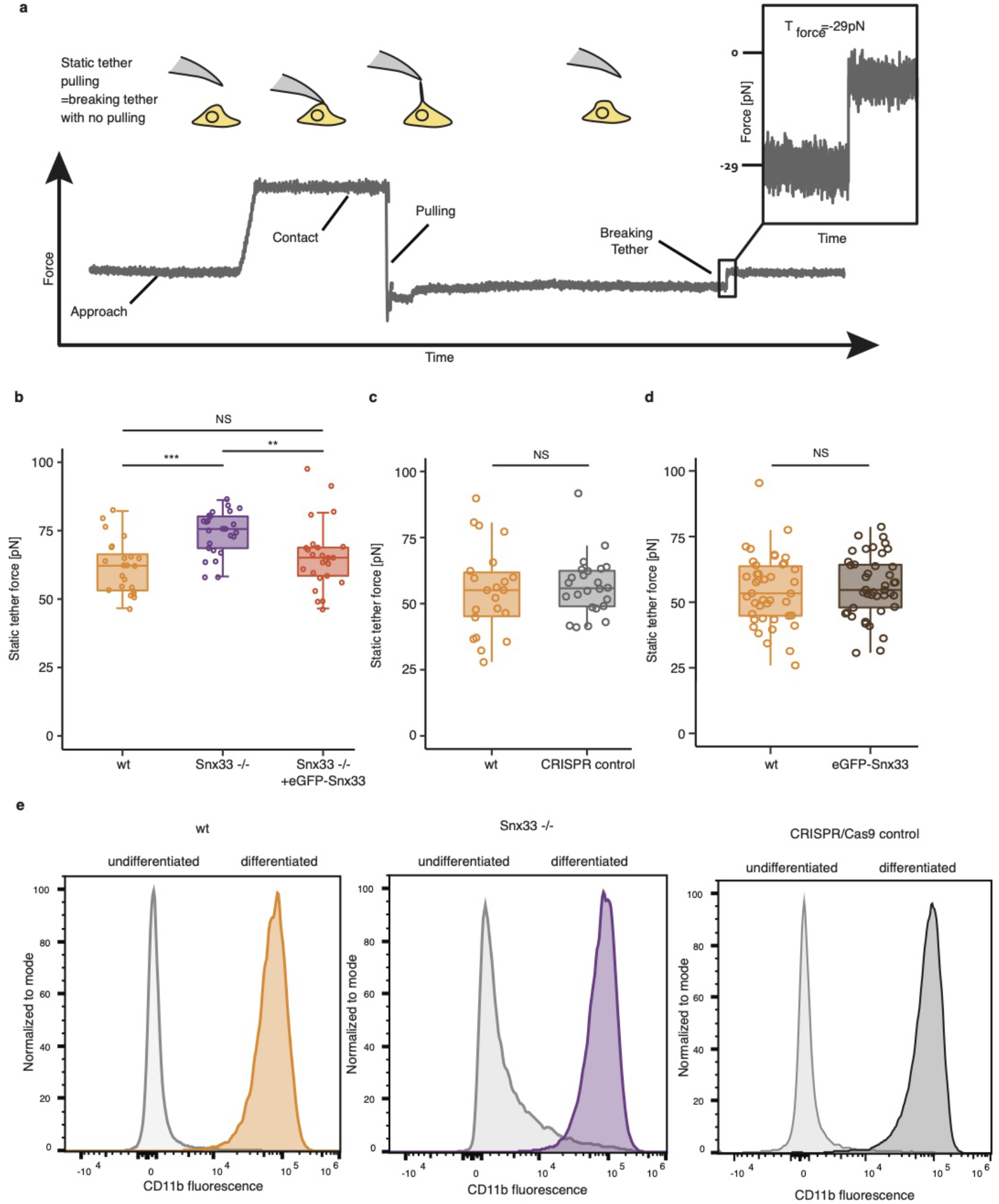
Snx33 knockout increases apparent membrane tension. **a)** Schematic of static tether pulling experiments. **b)** Mean static tether force of wt (n = 24), Snx33-/- (n = 26) and Snx33-/- with overexpressed eGFP-Snx33 (n = 25) from 3 independent experiments. **c)** Mean static tether force of wt (n = 27) and CRISPR control (n = 28) cells from 4 independent experiments. **d)** Mean static tether force of wt (n = 42) and eGFP-Snx33 over-expressed (n = 42) cells from 6 independent experiments. Overexpression of Snx33-GFP on the wt background did not decrease the tether force, suggesting that a gain of function is not sufficient to alter the leading edge or its effects on membrane mechanics. **e)** Representative histograms of CD11b intensity of wild-type, Snx33-/- and CRISPR/Cas9 control of HL-60 cells before and after 5 days of differentiation. Statistics: t-test and Mann-Whitney-U-test. p<0.001 (***), p<0.01 (**), p<0.05 (*).

**Supplementary Figure 7:**
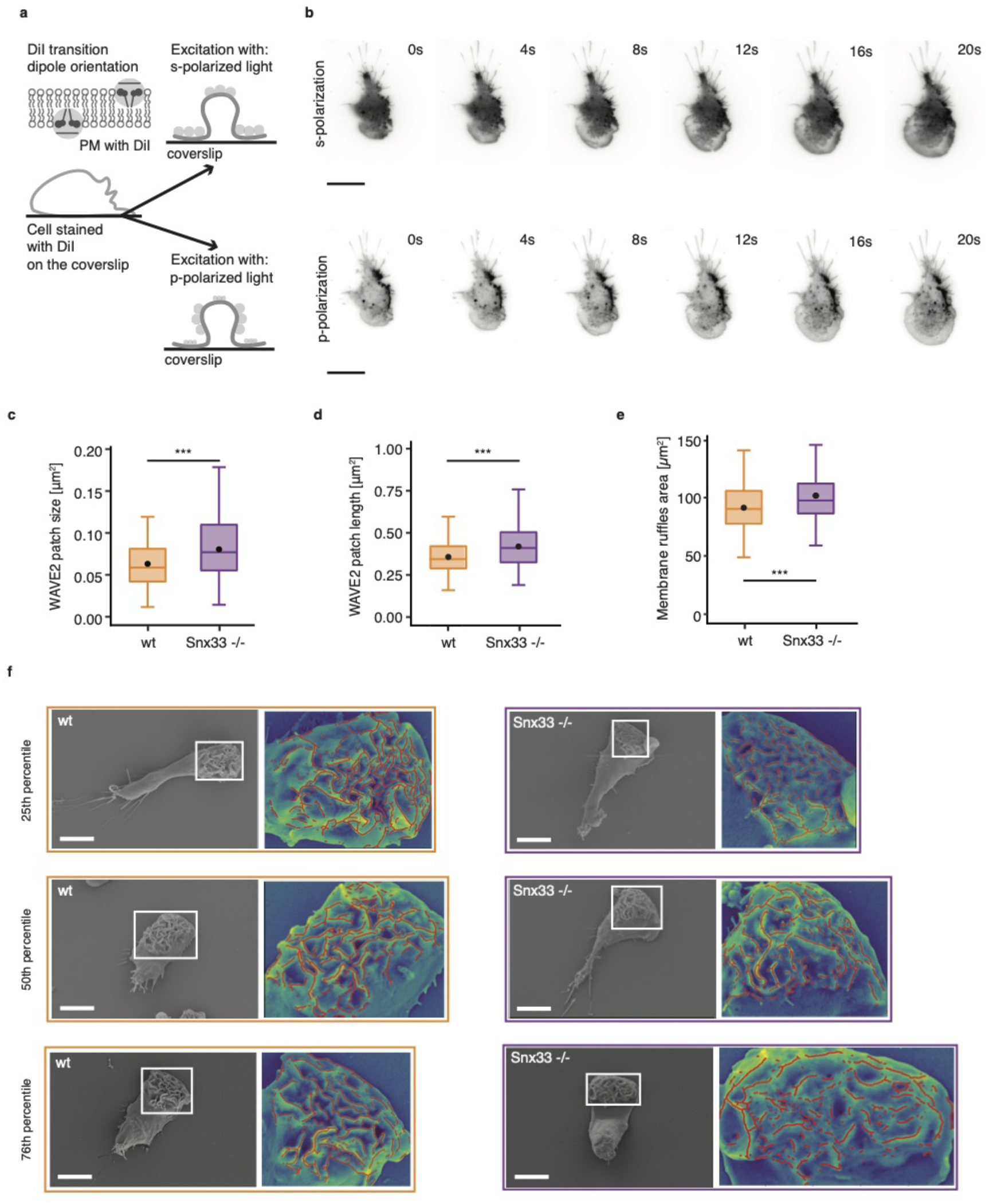
Membrane topography imaged by pTIRFM. WAVE2 pattern size and ruffle wavelength increases in Snx33-/- cells. **a)** Schematic illustrating the principles of pTIRFM imaging using carbocyanine dyes (*DiI*). **b)** Time lapse pTIRFM imaging of a cell using s-polarization (upper panel) and p-polarization (lower panel). Quantification of **c)** WAVE2 patch size and **d)** length in wt and Snx33-/- cells imaged by TIRFM. n=82 (wt), n=78 (Snx33-/-). **e)** Quantification of membrane ruffles area in wt and Snx33-/- cells imaged by SEM. n=175 (wt), n=170 (Snx33-/-). **f)** SEM images (25^th^, 50^th^ and 76^th^ percentile) with zoom-in of the overlay of the leading-edge and ruffle segmentation (red) for wt and Snx33-/- cells. Statistics: t-test or nonparametric Mann-Whitney-U-test. Scale bars = 10 μm. p<0.001 (***), p<0.01 (**), p<0.05 (*).

**Supplementary Figure 8:**
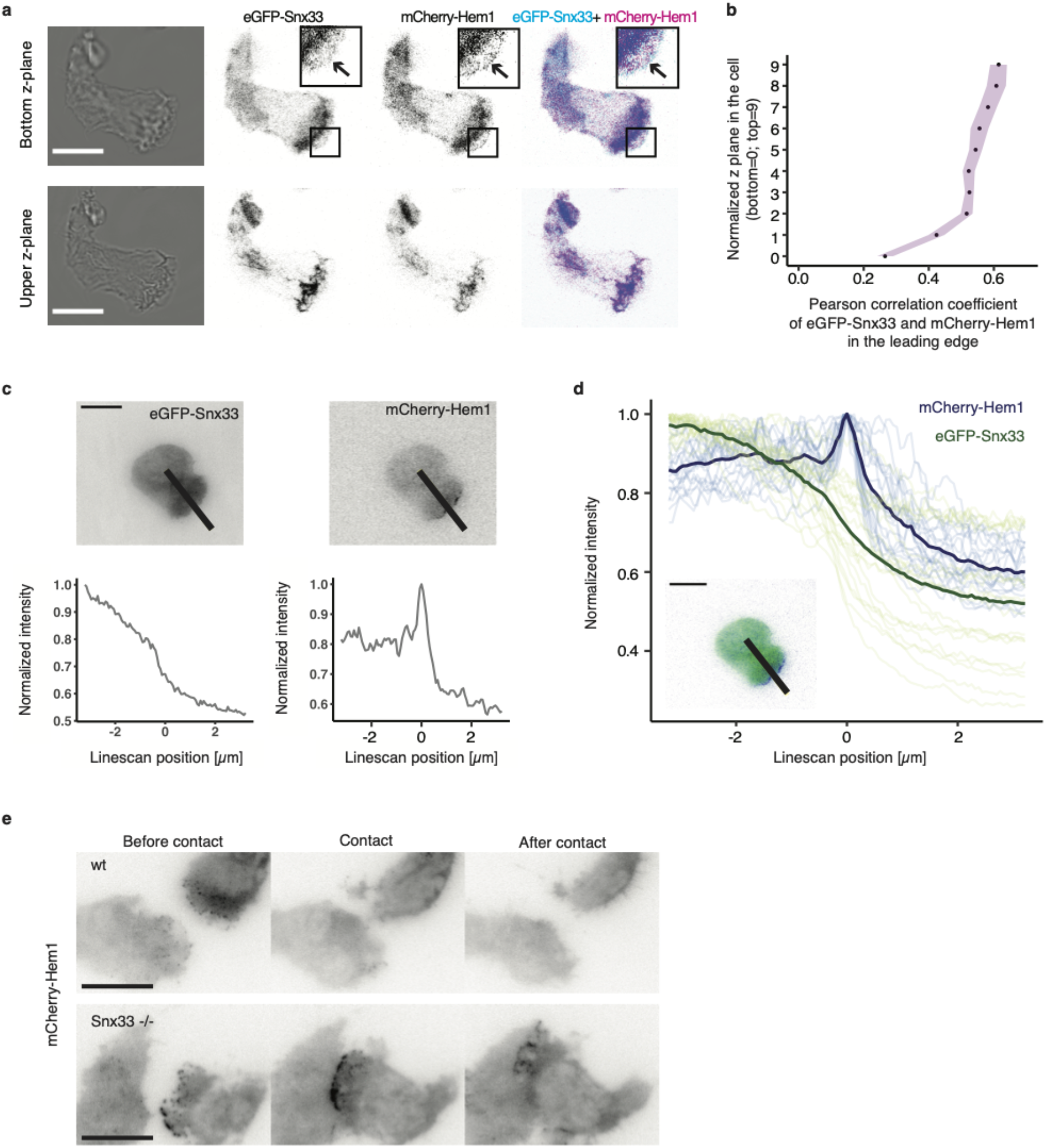
Snx33 is excluded from the protruding edge. **a)** Bright-field and confocal images of a cell, Snx33 and WAVE2 in the bottom and upper z-planes. Arrows indicate the most protruding edge at the cell bottom. **b)** Pearson correlation of Snx33 and WAVE2 in normalized z-planes in the leading edge. n=10. Purple regions denote standard error of the mean. **c)** Exemplary images of fluorescently-tagged Snx33 and WAVE2 and line scans through the leading edge in cells using TIRFM. **d)** Snx33 and WAVE2 normalized intensity at the protrusion edge. **e)** TIRFM images of WAVE2 signal before, during and after the contact event in wt and Snx33-/- cells. Scale bar= 10 μm.

## Acknowledgements

We thank Jan Ellenberg for critical feedback on the manuscript, Nir Gov and Ulrich Schwarz for fruitful discussions, and the Life Science Editors for editing assistance. The plasmid with hSnx33 was a kind gift from Duanqing Pei. We thank Brian Graziano for providing protocols, reagents and key advice to generate CRISPR knockout HL-60 cells. We thank Jakub Czuchnowski for advice on image analysis. We thank the EMBL flow cytometry core facility, the EMBL advanced light microscopy facility and the EMBL genomics core facility for support and advice. We thank the EMBL genome biology computational support (and specially Charles Girardot and Jelle Scholtalbers) for critical assistance during RNAseq analysis. We acknowledge the financial support of the European Molecular Biology Laboratory (EMBL) to A.D-M., Y.S. and A.E., the EMBL Interdisciplinary Postdocs (EIPOD) fellowship under Marie Sklodowska-Curie Actions COFUND to M.S.B., the BEST program funding by FCT (SFRH/BEST/150300/2019) to S.D.A. and the Joachim Herz Stiftung Add-on Fellowship for Interdisciplinary Science to E.S.

## Author Contributions

A.D.M., and E.S conceived the project and designed the experiments. E.S., M.B. and J.S., performed the experiments with EM advice from Y.S and microfluidic support from M.S.. E.S. and S.D.A. analyzed the data with support from A.E. and A.K.. A.E. performed the theoretical analysis. A.D.M., E.S and A.E wrote the manuscript. All authors contributed to the interpretation of the data, read and approved the final manuscript.

## Methods

### Cell culture

HL-60 cells were grown in RPMI 1640 media with 10% heat-inactivated FBS (#10500-064, Gibco) and 1% Penicillin-Streptomycin (#15140-122, Gibco) in a humidified incubator at 37°C with 5% CO_2_. Cells were differentiated by adding 1.5% DMSO (#D2438, Sigma Aldrich) and used after 5 days. Each independently-differentiated batch was treated as a biological replicate. For starvation, cells were kept for 1 hour in FBS-free RPMI 1640 media with 0.3% fatty acid free BSA (#A7030-10G, Sigma Aldrich). For imaging or fixation, differentiated HL-60 (dHL-60) cells were plated on fibronectin-coated (0.01mg/ml, #356008, Corning) glass-bottom dishes (#627860, Greiner bio-one) and allowed to adhere for 10 minutes in growth media. Next, cells were washed and stimulated with 10 nM fMLP (#F3506-5MG, Sigma Aldrich). For lowering adhesion, the coating was supplemented with 5% molar BSA (#A7030-10G, Sigma Aldrich). To generate stable cell lines with GFP-Snx33, mCherry-Hem1 (component of the WAVE2 complex, referred to as WAVE2) and mCherry-CAAX lentiviral transduction was used as described previously^33^. Cells were sorted on a BD FACS Aria™at EMBL Flow Cytometry Core Facility.

### Generation of knockout cell line by CRISPR/Cas9

CRISPR/Cas9 generation in HL-60 cells was performed as described previously^34^. Cloning of the target guide sequence to target *Snx33* was performed as previously described^45,46^ (Forward: *CACCGctgggacgacGGATGCACAG*; Reverse: *aaacCTGTGCATCCgtcgtcccagC*). Cells expressing BFP-tagged Cas9 were single-cell sorted in 96-well plate on BD FACS Aria™ Fusion at EMBL Flow Cytometry Core Facility. Single-cell clones were verified by genomic DNA amplification by Touchdown PCR^47^ and sequencing, followed by Western blot of selected clonal lines.

### Western blot

For immunodetection of Snx33 and GAPDH, 6×10^6^-1.2×10^7^ HL-60 cells were lysed in RIPA Lysis and Extraction buffer (#89900, Thermo ScientificTM) according to manufacturer’s instruction with supplementation of the protease inhibitors (#4693159001, Roche). Samples were denatured with 4xLaemmli buffer (#161-0747, BioRad) containing 10% β-mercaptoethanol (#m6250, Sigma Aldrich) at 95°for 5 min. After SDS-PAGE and transfer, PVDF membrane with transferred proteins was blocked in 5% BSA in TBST and incubated over-night with 1:1000 dilution of anti-Snx33 (#orb331346, Biorbyt) or 1:80 000 dilution of anti-GAPDH (#NB300-221, Novus Biologicals). The blot was developed with secondary antibodies at 1:10 000 dilution of Donkey-Anti-Rabbit-HRP (711-035-152, Jackson Immuno Research) or Goat-Anti-Mouse-HRP (115-035-062, Jackson ImmunoResearch).

### CD11b staining of HL-60 cells

After starvation, 1×10^5^ of undifferentiated and differentiated HL-60 cells were stained with Anti-Hu CD11b Alexa Fluor® 488 antibody solution (#A4-681-T100, Exbio). Fluorescence was measured on a Cytek® Aurora (Cytek) at the EMBL Flow Cytometry Core Facility and further analyzed and plotted using FlowJo.

### RNA sequencing

Total RNA samples obtained from 3 biological replicates were purified using RNeasy Mini Kit (#74104, Qiagen) according the manufacturer instructions with a DNase digestion step (#79254, Qiagen). To ensure high quality, samples were analyzed on an Agilent 2100 Bioanalyzer (Agilent Technologies). RNA sequencing was performed on an Illumina NextSeq 500 platform as NextSeqHigh-75 SE at the EMBL genomics core facility. For sequence alignment the hg19 reference genome was used. Differential expression analysis was performed with a custom-made Galaxy pipeline using a DESeq2 package.

### Imaging

TIRFM images of live cells were acquired on a Nikon Ti Eclipse inverted microscope with a TIRF objective CFI Plan Apo Lambda 100x Oil (#MRD01905, Nikon) and sCMOS camera controlled by NIS-Elements (Nikon). Sample drift was reduced using an autofocus system (Perfect Focus, Nikon) for time lapse imaging.

Confocal images of fixed cells were obtained with a silicone objective UPLSAPO 60X S (NA 1.3; WD 0.3 mm) on an Olympus FV3000 inverted microscope at EMBL advanced light microscopy facility.

Epifluorescent and bright-field imaging of fixed cells was performed using a 40x objective (#MRD00405, Nikon), the SOLA SE II and 100W halogen lamps (Nikon) using appropriate filter sets.

Polarized TIRFM (pTIRFM) modality was implemented based on previous work^48–53^. For imaging, dHL-60 cells were stained before plating with carbocyanine dye DiI (#D3911, ThermoFisher Scientific).

### Fixation and F-actin staining

Migrating dHL-60 cells were fixed by adding fixation buffer (2x) to growth media (1:1 v/v) and incubated at 4°C for 1 hour. Fixation buffer (1x) contains 3.7% paraformaldehyde (#28908, Thermo Scientific), 1x intracellular buffer (140 mM KCL, 1 mM MgCl_2_, 2 mM EGTA, 20 mM HEPES, pH 7.5), 320 mM sucrose (#S0389-500G, Sigma Aldrich) and 0.2% BSA (#A7030-10G, Sigma Aldrich). Cells were washed and stored in dPBS. For permeabilization and staining, cells were re-suspended in intracellular buffer (1x) containing 0.2% of Triton X-100 (#T8787, Sigma Aldrich). If applicable, phalloidin coupled with TRITC (#P1951, Sigma Aldrich) was added at this step. After washing with dPBS, cells were stored at 4°C in the dark or used immediately for imaging.

### Theoretical calculations

Calculations for Fig. 2j and Fig. 3a were performed in Wolfram Mathematica v12.1.1.0. For the details of the derivations please see the **Supplementary Note**.

### Cell migration assays in PDMS-based devices

PDMS-based microfluidic devices were prepared as previously described^17,54,55^. The devices used for migration of dHL-60 cells had heights of 2.8 μm and 3.13 μm for channels with decision point and channels with constriction, respectively. The decision channels had constrictions of 2, 3, 4 and 5 μm in two arrangements. The channels with single constrictions were 2 μm. To visualize nuclei and cell body, Hoechst 33342 (#62249, Thermo Fisher Scientific) and TAMRA (Invitrogen) were added before the introduction of cells into the PDMS device. Cell migration towards chemoattractant (fMLP) was imaged on an inverted wide-field Nikon Eclipse microscope using 20x/0.5 PH1 air objective, equipped with a Lumencor light source (390 nm, 475 nm, 542/575 nm), an incubation chamber and the heated stage with CO_2_. The acquired data were analyzed using ImageJ software and manually curated. Only single cells that moved through the entire channel were considered for analysis. All the parameters were quantified based on the nuclei signal.

### Tether extrusion using atomic force spectroscopy

Apparent membrane tension was measured by extruding plasma membrane tethers. For measurements, Olympus BioLevers (k = 60 pN/nm) from Bruker were mounted on a CellHesion 200 AFM (Bruker), which is integrated into an Eclipse Ti inverted light microscope (Nikon). Cantilevers were calibrated using the thermal noise method and coated with 2.5 mg/ml Concanavalin A (#C5275, Sigma Aldrich). Prior to the measurements, cantilevers were rinsed in dPBS. For tether measurement, the cantilever was position over the cell, preferably over the leading edge. Measurements parameters for static tether pulling experiments were as followed: approach velocity was set to 1 μm/s, contact force to 100–300 pN, contact time to 5–10 s, and retraction speed to 10 μm/s. After a 10 μm tether was pulled, the cantilever position was held constant until it broke, but no longer than 30 s. In every experimental repetition the conditions’ order was randomized. For every cell at least 3 different tether measurements were taken.

The data analysis was performed using the JPK Data Processing Software. For assessing the magnitude of membrane tension based on tether force measurements, the following formula was used^33^:

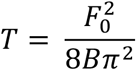

where *F_0_* is the tether force measured by the AFM and *B* is the bending rigidity of the plasma membrane, which we assume to be invariable between different experimental conditions (2.7×10^-19^ Nm based on previous measurements^56,57^).

### Image analysis

For confocal images, only the z-planes that contained the top 80% intensity of mCherry-CAAX (membrane marker) were considered based on line scans covering the entire resliced maximum intensity z projection. A channel of interest (ChoF1) was used for mask generation based on automatic Otsu segmentation. A custom-made ImageJ script allowed to calculate the Pearson correlation coefficient (PCC) for every z-plane of ChoF1 with ChoF2 based on the mask of ChoF1 using the in-built Coloc2 ImageJ plugin. Z-slices were assigned to 10 bins and the mean with standard error of the mean for every bin was calculated.

For epifluorescence images, single cells were manually selected using the ImageJ software. To segment the cell body, the contrast of bright-field images was enhanced using the equalize histogram function followed by a canny edge detection, a semi-manual closing of the obtained edges and filling the holes. To segment the leading edge, a similar strategy was used with semi-manual selection of the region with the enriched edges after the canny edge detection with higher values step. Based on the segmentation, a custom Python script was used for measurements. Eccentricity was calculated by ellipse fitting and further division of ellipse foci distance by major axis length.

For analysis of migrating cells imaged by TIRFM, a segmentation of the cell mask (based on mCherry-CAAX signal) and of the WAVE2 mask (based on eGFP-Hem1 signal) were acquired using the machine learning-based ilastik software^19^. Further image analysis was achieved using an in-house built program implemented in Python. The angle at which cells are moving was calculated based on the center of mass for 3 consecutive frames. The leading edge was defined as the difference between two consecutive frames where at least one pixel of WAVE2 mask is present per cluster. Leading edge length was defined as a number of pixels in the outside perimeter of the leading edge. For the analysis of cellcell contacts the same segmentation strategy was used to segment individual cells. Cell-cell contact was defined as the perimeters’ intersection of both cells.

For membrane topography analysis of SEM data, the leading edge area with ruffles was manually segmented on median filtered images. Next, ridges were detected within the segmented regions using the Meijering filter^58^. Ridges were later segmented using automatic Otsu thresholding and skeletonized using a custom Python script. Inversion of the number of pixels within the skeletonization per leading edge area corresponds to the effective the ruffle wavelength.

For analysis of signal enrichment at the cell edge based on TIRFM images, data on fluorescent intensity along the line of equal width and length were extracted from both channels of interest (eGFP-Snx33, mCherry-Hem1) using ImageJ. Data were aligned according to the highest fluorescence intensity of mCherry-Hem1 and normalized. Normalized mean intensity was calculated in both channels of interest for every point along the line scan and plotted using R.

### Scanning electron microscopy

After 30 min of 10 nM fMLP stimulation, cells were fixed in 2,5% GA (#16220, EMS) in 0,1M PHEM buffer by adding 37°C double strength fixative (5% GA in 0,1M PHEM) directly 1:1 to the cell medium. After 10 minutes incubation, the fixative was replaced by fresh single strength fixative and cells were further fixed at room temperature for 1h. After fixation, cells were washed 2 times in 0,1M PHEM and 2 times in 0,1M cacodylate buffer. Next, they were postfixed for 2h on ice in freshly prepared and filtered 1% OsO_4_ (#19190, EMS) and 0,8% potassium ferrocyanide (K_4_[Fe(CN)_6_]*3H_2_O, #4984, Merck) in 0,1M cacodylate buffer. After postfixation, the cells were washed 4 times in H_2_O, and left at 4°C until further processing.

Next, cells were treated with freshly prepared and filtered 1% tannic acid (TA, CAS#1401-55-4, EMS) in water using a Pelco BioWave microwave for seven 1-minute cycles alternating between 150 W and 0 W power. Steady temperature was set to 23°C and vacuum to on for all steps. After TA treatment, cells were washed 2x in H_2_O on the bench and 2x in H_2_O in the microwave for 40 s per step at 250 W power. Cells were then treated with 1% UA (#77870, Serva) in H_2_O using the same microwave program as for TA. After washing once in H_2_O and twice in 25% EtOH, cells were dehydrated in a graded series of ethanol (25% - 50% - 75% - 90% - 100% - 100%) using a microwave program with step length of 40 s and 250 W power, with a steady temperature at 4°C and without vacuum. Finally, the cells were infiltrated with a graded series of Hexamethyldisilizane (HMDS, CAS# 999-97-3, Sigma Aldrich) in ethanol (25% - 50% - 75% - 100% - 100%) using a microwave program with 6 steps of 1 minute each, with a power of 150 W for step 1, 3, 4 and 6, and 0 W for steps 2 and 5. After the final 100% HMDS infiltration, all HMDS was removed, and coverslips were left to dry overnight. Silica gel with moisture indicator (Merck) was added in 4 empty wells (corners) in the 24-well plate to remove excess humidity.

After drying, coverslips were mounted in aluminum stubs (Agar Scientific G301F) using carbon tape, and sputter coated with a layer of gold for 180 s at 30 mA current using a Quorum sputter coater model Q150RS.

Imaging was performed on a Zeiss Crossbean 540 microscope, using 5 kV acceleration voltage and 700 pA current for the electron beam, with a working distance of 5 mm. A secondary electron detector (SESI) was used for signal detection, and all images were acquired with a pixel size of 28,9 nm/pixel.

### Statistical analysis

Statistical analyses were performed using R, while data visualization by both R and Adobe Illustrator^®^. Normality of data distribution was tested by Shapiro-Wilk test. Two-tailed t-test was used for normal distribution. Otherwise, a non-parametric Mann-Whitney-U-test was used, if not indicated differently. In all box-plots, the lower and upper hinges correspond to the first and third quartiles (the 25th and 75th percentiles). The upper whisker extends from the hinge to the largest value, but no further than 1.5*IQR (distance between the first and third quartiles). The lower whisker extends from the hinge to the smallest value, but no lower than 1.5*IQR of the hinge. Data beyond the end of the whiskers are plotted as black dots. Black line and dot correspond to the median and mean, respectively.

## Supplementary Note: Curvature-dependent mechanochemical patterning

Many migrating cells exhibit characteristic patterns of actin activity and membrane curvature at their front [1]. These have been studied in the past by considering the complex interplay of various actin-regulatory molecules [7, 8]. Interactions with the curvature of the plasma membrane have also been investigated, in particular the effect of curvature-sensing and inducing proteins that promote the polymerization of actin [4, 5, 9–11]. In immune-like HL-60 cells, the BAR domain containing protein Snx33 is upregulated during migration. Based on its structure, Snx33 is likely to bind membrane regions with weak inward curvatures, and we find that disrupting Snx33 impacts the cell-scale organization of the actin cytoskeleton. We investigate here a putative role of Snx33 on the patterning capacity of the leading edge. We consider a minimal set of equations to describe the dynamics of membrane curvature coupled to the concentration field of an activator of actin polymerization. We analyze the mechanochemical instabilities present in the system, and delineate the parameter region in which spontaneous patterning can occur.

### 1 Main equations

In the following, we derive the coupled system of equations consisting of a curvature-dependent reaction-diffusion equation for an activator of actin polymerization, and the equation for the curvature of the membrane.

#### 1.1 Activator equation

We consider the concentration field *A* of a generic activator of actin such as WAVE2. Because actin protrusions at the cell front are often thin structures with a dense meshwork and an average filament orientation towards the leading edge [13], we neglect variations of *A* across the thickness of the protrusion and assume that diffusive transport of actin regulatory molecules occurs predominantly in the direction of average filament alignment *x*. We consider self-activation of *A* with a characteristic time constant *τ*_A_ and introduce the curvature-dependent regulatory term *r*_C_(*C*). From the continuity equation for the surface concentration of activator molecules, we then obtain the following partial differential equation

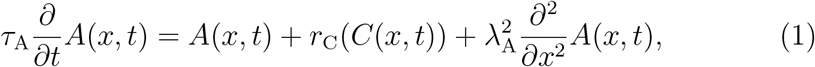

in which 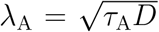 denotes a reaction-diffusion length scale, with *D* the diffusion coefficient along the cell migration axis *x*.

#### 1.2 Curvature equation

The curvature *C* depends on the mechanical properties of the plasma membrane. We consider a Helfrich membrane without spontaneous curvature with a bending rigidity *κ* and surface tension *γ* [2, 6]. In the weak bending approximation and with periodic boundary conditions, the conservation of momentum leads to the following equation for the curvature profile along *x* [3, 12]

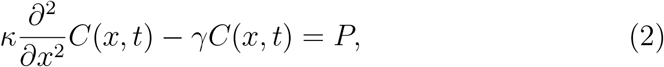

in which *P* = *P*_ext_ − *P*_int_ denotes the pressure difference between the outside and the inside of the cell. We assume that actin polymerization generates an active pressure field which acts on the membrane from within the cell that depends on *A*(*x, t*), and we neglect any pressure difference in the absence of activator molecules. With this term and introducing the membrane persistence length 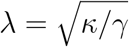, Eq. 2 reads

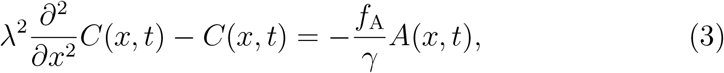

in which we denote with *f*_A_ the force per activator molecule.

### 2 Regulation in the limit of small curvatures

Generally, the shape of the regulatory function *r*_C_(*C*) is unknown and depends on the properties of the curvature-sensing proteins involved. The shape of the dominant BAR protein in our experimental system suggests that curvature sensing is relevant in the regime of small curvatures *C*(*x, t*) = *δC*(*x, t*) (**Supplementary Fig. 2**). Assuming differentiability of *r*_C_ with respect to *C*, we may write a general expansion of the form

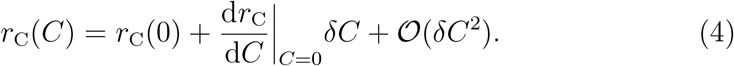

The magnitude of the linear coefficient depends on the activity of curvature-dependent regulatory proteins. In the following, we assume that no curvature regulation takes place for a completely flat membrane sheet, i.e. *r*_C_(0) = 0.

### 3 Patterning regime

We now explore the consequences of curvature-regulatory coupling between membrane shape and actin activity by analyzing the stability of the steady state *C* = 0 and *A* = 0, in which the membrane is flat and no actin activity is present. Introducing rescaled variables and the normalized curvature coupling coefficient

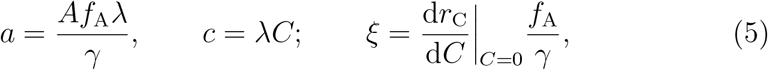

we may write Eqs. 1 and 3 in dimensionless form as

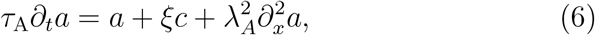

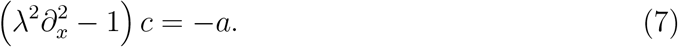

A small perturbation away from the uniform steady state can be written as

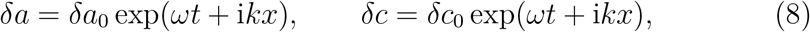

in which *k* is the perturbation wavenumber and *ω* the corresponding growth rate. Substituting Eq. 8 into Eqs. 6 and 7 and keeping terms to linear order in the perturbation, we obtain the dispersion relation

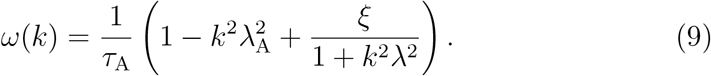

The stability diagram as a function of the length scale ratio *λ*/*λ*_A_ and the curvature coupling constant *ξ* is shown in **Fig. 2j**. For positive coupling, the growth rate is positive and maximal at *k* = 0, describing a uniformly advancing flat sheet in the linearized system. For *ξ* < 0, there is a region in which spontaneous patterning occurs, i.e. where *ω*(*k*_Max_) > 0 with *k*_Max_ > 0. Two types of instabilities are present in the system. For *λ* < *λ*_A_, stable and unstable regions are separated by a type-III instability at the critical parameter value 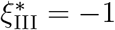, with timescale *τ*_A_ and coherence length 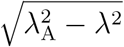. For *λ* > *λ*_A_, the transition between stable and unstable regions corresponds to a type-I instability, and the critical line is given by

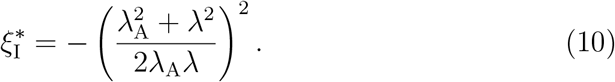

Here, the characteristic timescale is 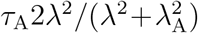 and the coherence length is 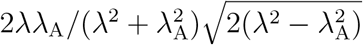. The pattern wavelength at the maximum is given by

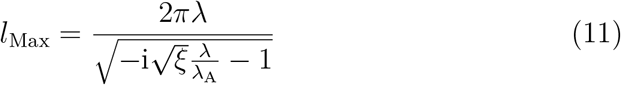

and shown in **Fig. 3a**. Thus, negative curvature feedback can lead to the spontaneous formation of patterns. The wavelength of the fastest-growing mode increases when the magnitude of the curvature coupling constant *ξ* decreases.

We note that by coupling the Snx33-mediated feedback circuit to a second positive feedback loop, for example driven by a second polymerization-promoting curvature sensing protein species as considered here [5], one may recover dynamical behaviors such as traveling waves, which are observed in migrating cells.

